# Loss of Cysteine Rich domain was critical for evolution of heterodimerization in Toll proteins

**DOI:** 10.1101/2021.01.18.427181

**Authors:** Poonam Ranga, Suresh Kumar Sawanth, Nirotpal Mrinal

## Abstract

Toll proteins play roles in immunity/development which have largely remained conserved. However, there are differences in Toll biology as mammalian TLRs recognise pathogen associated molecular patterns (PAMPs) but not their invertebrate homologues. The reason for the same is not known. One critical molecular difference is absence of Cysteine Rich Domain (CRD) in vertebrate Tolls and their presence in invertebrates. Interestingly, in Drosophila, all Toll proteins have CRD except Toll9. This provided us the appropriate model to investigate significance of loss of CRD in Toll evolution. CRDs nudge protein dimerization by forming disulphide bonds hence we asked if they did same in Drosophila Toll-proteins. We tested if, Toll-1(which forms homodimer) can heterodimerize with Toll-9. We found that wildtype Toll-1 didn’t interact with Toll9 however; when CRD of Toll1 was deleted/mutated it formed heterodimer with Toll9. This indicates that presence of CRD limits Toll proteins to form homodimer and thus its loss was a critical event which pushed Toll proteins towards heterodimerization. We further show that Drosophila Toll9 can directly bind dsRNA, a PAMP. Interestingly, dsRNA affinity for toll-9 monomer was twice as that for the dimer, which can be attributed to CRD loss. Thus, we show that loss of CRD was a major step in Toll evolution as it resulted in functional diversity and was a first step towards heterodimer formation. Therefore, we propose that CRD loss was under positive selection and also that heterodimerization of Toll-proteins is an evolved property.

**One line summary:** Loss of Cysteine Rich Domain in Drosophila Toll9 and recognition of dsRNA.

## Introduction

The Toll pathway plays a central role in immunity and development and is highly conserved across animal groups. Toll was discovered first in Drosophila as maternal gene of Dorsal group(1,2). Laterit was also discovered in mammals and named as TLRs(3). These proteins are highly conserved in structure and function from early metazoan, Nematostella to insects to vertebrates (4, 5). Recently a toll resembling protein (SDTLR) has been identified in Porifera-*Suberitesdomunculawhich* indicatesits early origin(6–8). SDTLR is primitive as it only has a TIR domain and a transmembrane anchor but no Ig and LRR domains. This suggests that LRR which provides seat for ligand binding in Toll proteins was absent in primitive Toll (8). All LRR proteins form horseshoe shaped structure with a concave and a convex surface(9). The inner concave face of the horseshoe shape has LxxLxLxxN repeats. It is believed that extraordinary ability of these proteins to form dimers and recognize diverse variety of ligands from hydrophobic to hydrophilic molecules is because of the structural patterns in the horseshoe shaped dimeric structures(10). It is to be noted that formation of TLR homomultimers or heteromultimers does not require ligand binding but these homo/heterodimers are not functional till a ligand binds (11–13). This characteristic horseshoe shaped structure of Toll dimers is stabilized by cysteine disulphide bridges at the N and C terminal ends of the LRR domain(14). Sequence analysis suggests that all Toll proteins surely have one cysteine cluster called as Cystine rich domain (CRD) sandwiched between transmembrane domain and the LRR domain(15). The second CRD, present within the LRR domain(CRD-LRR), is absent in all chordates but present in invertebrates *viz*. insects, annelids, molluscs etc.(16) (Figure 1). Drosophila is an exception among lower organisms as it has both types of toll proteins. Toll9 of Drosophila lacks the CRD within the LRR domain and thus resembles mammalian TLRs in structure while rest of Drosophila Toll proteins have two CRDs like any insect toll(*17*).

**Fig 1.**
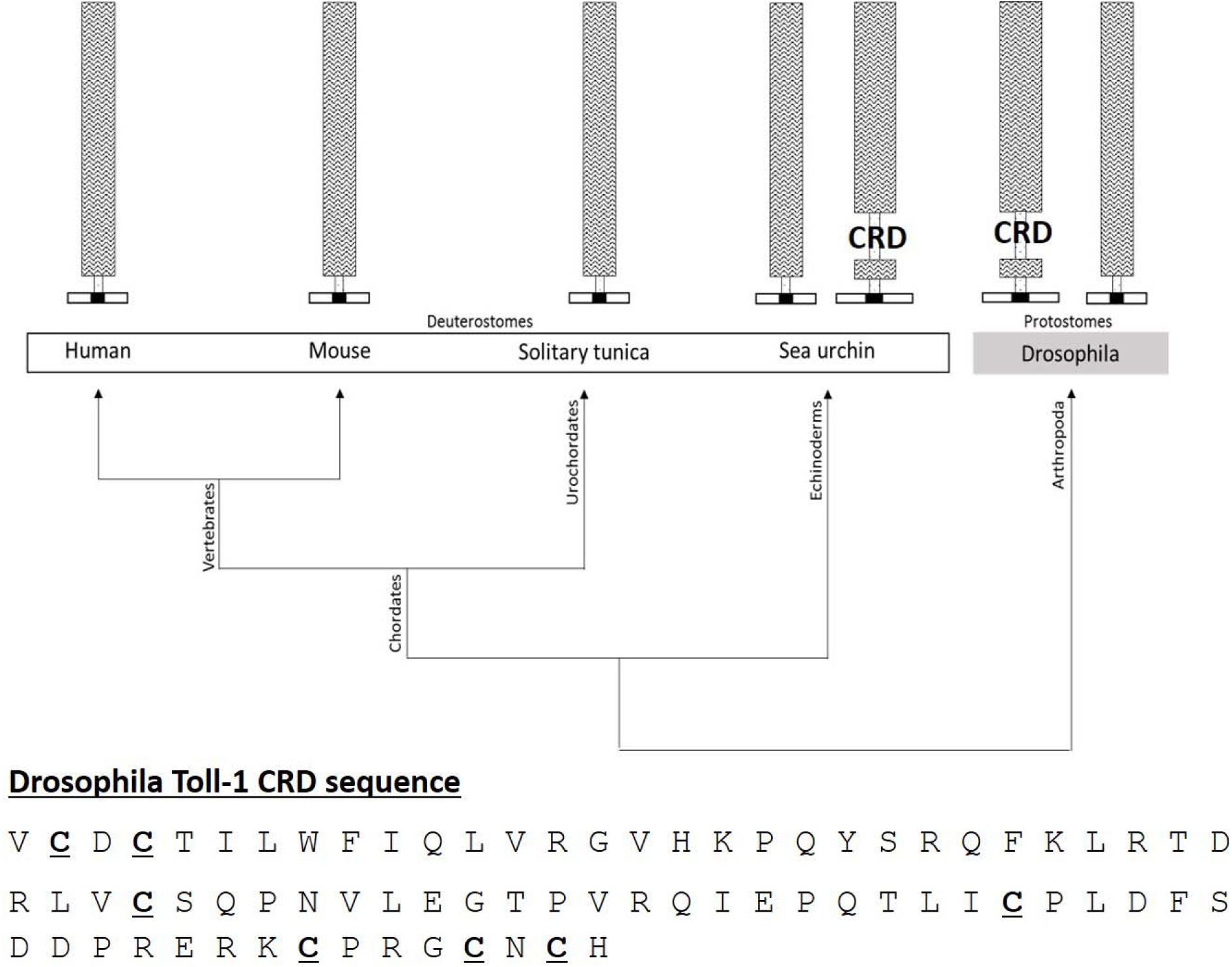
Evolution of Toll proteins. Phylogenetic tree shows loss of CRD over the course of evolution of vertebrates. Invertebrates largely have Toll with CRD with the exception of *Drosophila* where Toll-9 lacks LRR-CRD. Seven cysteine residues in the CRD are highlighted.

At the core of Toll biology is their ability to form homo/heterodimers within the family. While mammalian TLRs form homodimers as well as heterodimers, insect tolls are known to form homodimers only(*18–21*). A closer analysis of LRRs of insect and mammalian Toll reveals loss of CRD-LRR in mammals as the most noteworthy difference. CRDs play many roles including protein-protein interaction hence their presence and absence usually has strong phenotypes. e.g. Loss of CRD in dystrophin gene leads to Duchenne muscular dystrophy (DMD) while deletion of the same in TNF receptor abolishes TNF binding (*22, 23*). Also, the first 2 CRDs in Sog protein of *Drosophila* are accountable for BMP activity (*24*).Clearly, they play important roles.

*Drosophila* Toll1 possesses a cysteine rich domain between 17^th^ and 18^th^ LRR. The LRR of Toll1 interacts with LRR of second monomer of Toll1 to form a dimer (14). This dimer then binds with spätzle to initiate toll signalling (14, 25). However, there is no report to suggest binding of Toll1 with other toll paralogs not even Toll2 (18wheeler) which is co-expressed with Toll1 upon Gram positive bacteria infection. This suggests that the LRRs of Toll1 and 18w don’t interact. Our aim was to understand role of CRD within the LRR domain in Toll dimerization. Drosophila has eight insect type Toll proteins (those with two CRDs) and one mammalian type Toll protein, Toll-9 (with only one CRD). Presence of a naturally occurring Toll protein without LRR-CRD (Toll9) made *Drosophila* an ideal model to explore the role of CRD in Toll function.

Here, we show that LRR-CRD was required for formation of Toll homodimers and an impediment in Toll heterodimerization. Since, heterodimerization added to functional diversity and hence was evolutionary selected. As a result LRR-CRD experienced negative selection and hence was lost during the course of evolution. Thus, heterodimerization of Toll proteins is a gain of function phenotype resulting from loss of CRD.

## Results

While LRRs of Toll proteins have been extensively studied, the role of CRDs in Toll proteins remains unexplored. Here, we sought to dissect the purpose of CRD in Toll function. Since LRRs are sufficient for dimer formation then what purpose dose CRD serve?

### 1. CRD is required for formation of Toll1 homodimer

Towards this we created *toll*Δ*CRD* construct where the LRR-CRD was deleted in *toll-1* (Fig. S1). Next, we tested if *Toll9* can interact with Toll-1 by forming a Toll1-Toll9 heterodimer. Yeast two hybrid assay showed that wild type *toll1* did not interact with *toll9* at all (Fig. 2A). However, Toll1 deletion construct, *toll*Δ*CRD*. formed genetic interaction with *toll9* in Y2H assay (Fig. 2A). Because of deletion of CRD-LRR, *toll*Δ*CRD* construct was equivalent to *toll9* as both lacked LRR-CRD. This indicated that CRD of Toll1 inhibited interaction with Toll9 while deletion of the same resulted in dimerization with Toll9 (Fig. 2A). This supports our primary hypothesis that toll proteins with CRD form homodimers and those lacking CRD can form heterodimers.

**Fig 2.**
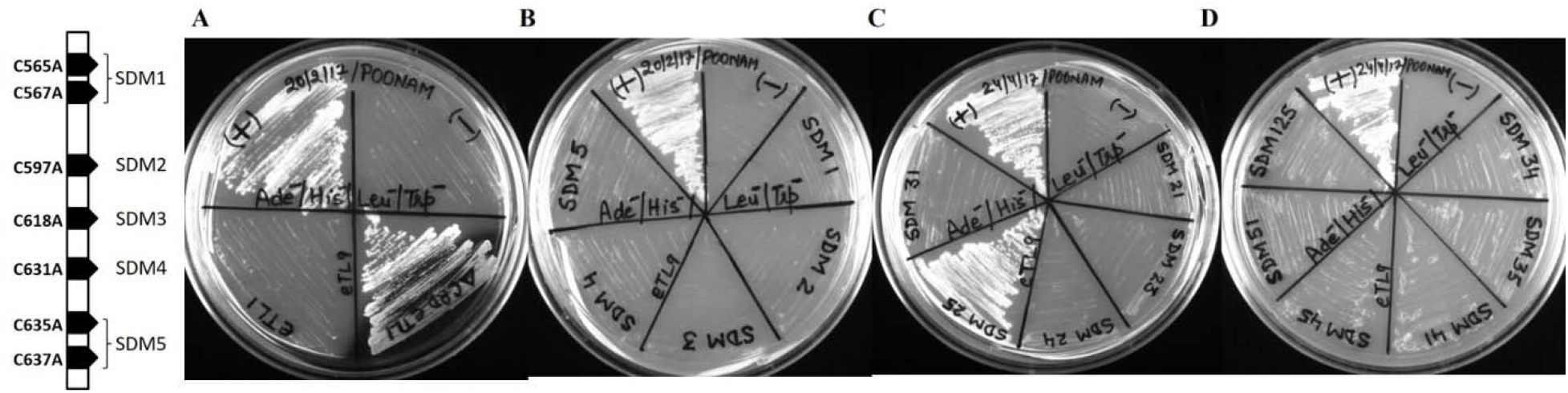
Yeast two hybrid assay to check interactions between toll1 and toll9. A) Positive interaction of toll9was seen with *toll*Δ*CRD* but not *toll1*. (B) SDM of individual Cys did not result in interaction with toll9. (C, D) SDM 25 formed heterodimer with *toll9*. Other mutant combinations did not show interaction with toll9. Absence of interaction between SDM 125 and toll9 was interesting and highlighted epistatic role of Cys 565 and 567. P53 and large T-antigen was used as positive control labeled (+) and mating between AD and BD empty vectors was used as negative control and labeled as (-). Leftmost panel shows Cys mutations.

### 2. Cysteines within the CDR form intra- and inter-molecular disulfide bonds to regulate dimer formation

After ascertaining that LRR-CRD played role in Toll homodimerization and inhibited heterodimerization we wanted to investigate the role of cysteine residues within the LRR-CRD of Toll1 in dimer formation (Fig. 1). CRD-LRR contains seven cysteine residues and using site directed mutagenesis we mutated each of them to Alanine to ascertain their role in dimer formation (Fig. 1).Cysteine residues at positions 565&567 (SDM 1), 597 (SDM 2), 618 (SDM3), 631 (SDM 4), 635&637 (SDM 5) were mutated to generate five mutant groups (Fig. 2, S2). We found that none of these *toll1* mutants resulted in interaction with *toll9* (Fig. 2B). The fact that these individual cysteine mutants did not lead to heterodimerization with toll-9 implied that multiple interactions were involved in Toll-1 dimerization. To test this possibility we generated 16 different combinations of mutants from these five SDM mutants (Fig. 2, Table-1).

**Table 1.** Summary of protein-protein interactions between toll mutants created by site directed mutagenesis (SDM) and toll9. (Y) indicates positive interaction while (x) indicates negative interaction or lack of interaction. Only SDM 25 shows heterodimer formation with Toll9. This table also summarises the epistatic relationship between different mutants. Most interesting epistatic interaction is shown by SDM1. Whenever SDM1 is used in any combination the heterodimer is never formed. This indicates that DSM1 is epistatic to all mutants tested here and has a role in heterodimer formation. Cys 567 is involved in intramolecular disulfide bond formation with Cys618 and may be important in maintaining the correct configuration for dimer formation. Cys565 supports heterodimer formation but how is not clear. But one can safely deduce that it is not involved in intermolecular disupfide bond formation. Epsistatic role of SDM1 is best explained by SDM125. SDM25 is the only mutant which favours heterodimer formation but once SDM muattions are added into this it does not show interaction with Toll9 (Fig. 2D).

| *Mutant* | *Mutation (C→A)* | *Heterodimer*<br>*with Toll-9* | *567-618* | *631-637* |
| --- | --- | --- | --- | --- |
| 1 | 565, 567 | x | x | Y |
| 2 | 597 | x | Y | Y |
| 3 | 618 | x | x | Y |
| 4 | 631 | x | Y | x |
| 5 | 635, 637 | x | Y | x |
| 21 | 565, 567, 597 | x | x | Y |
| 31 | 565, 567, 618 | x | x | Y |
| 41 | 565, 567, 631 | x | x | x |
| 51 | 565, 567, 635, 637 | x | x | x |
| 23 | 597, 618 | x | x | Y |
| 24 | 597, 631 | x | Y | x |
| 25 | 597, 635, 637 | Y | Y | x |
| 34 | 618, 631 | x | x | x |
| 35 | 618, 635, 637 | x | x | x |
| 45 | 631, 635, 637 | x | x | x |
| 125 | 565, 567, 597, 635, 637 | x | x | x |

Out of these 16 combinations that we tested, only one mutant combination SDM25 (generated by combining SDM2 and SDM5 mutants) comprising of Cys mutations at positions 597 (SDM 2), 635&637 (SDM5), resulted in heterodimer formation with Toll9 (Fig. S3, Table-1). Next, we added SDM1 mutations (Cys 565 and 567) to SDM25 to generate SDM125 which has 5 out of 7 Cys residues mutated (table 1). Interestingly, SDM125 did not form heterodimer with Toll9. This was surprising because we had hoped that SDM 125 would also show interaction with Toll9. Our assumption was that SDM25 mutations are the minimum required for interaction with Toll9 and any mutation in addition to this should only help the phenotype (interaction with Toll9). Clearly SDM1 proved epistatic to SDM25. For further understanding of epistatic relation between different mutants we mapped these mutations to Toll-1 structure.

The crystal structure of Toll1 revealed intramolecular disulfide bonds between Cys567-Cys618 and Cys631-Cys637 (14). So, these four Cysteines at positions 567, 618, 631 and 637 may not have a role in intermolecular disulfide bond formation and hence may not be critical for dimerization of Toll1. We inferred status of these two imtra-molecular disulfide bonds in all 16 mutants as shown in Table 1. It is apparent from this analysis that any mutation that disrupts the 2^nd^ intramolecular disulfide bond Cys631-Cys637 does not affect hetero-dimerization. Molecular interactions in mutants SDM25 and SDM125 are similar except for the formation of 1^st^ intramolecular disulfide bond between Cys567 and Cys618 (Table 1). It is clear that mutations at positions Cys565 and 567 are detrimental for interaction with Toll9 and hence are essential for heterodimerization. On the other hand Cys at positions 597 and 635 disfavour heterodimerization (table 1). The fact that SDM25 and *toll*Δ*CRD* construct have same phenotype suggests that Cys 597 and 635 are critical for dimer formation. While Cys at 597 and 635 favour homodimer formation, mutation at those positions favour heterodimer formation. Thus, we propose that Cys at 597 and 635 regulate Toll dimerization by forming intermolecular disulfide bonds (Fig. 2C, 2D and 3; Table 1). When these residues are mutated then intermolecular disulfide bonds are not formed which may nudge formation of heterodimer with Toll9 (Fig. 3). Consequently, Cysteine at 565 and 567 are essential and epistatic over Cys 597, 635 and 637 for heterodimerization. Contrarily, one can also say that Cys 597 and 635 residues are critical for homodimer formation in Toll1 and mutation at these positions probably leads to heterodimerization with toll9 (Fig. 2C). Cysteines in CRD form intra- and inter-molecular disulphide bonds thereby, making the region more rigid. Our genetic analysis gave us a pick into the roles of different cysteines of LRR-CRD which is in consonance with the structural data and clearly explains roles of different Cysteines in dimer formation at molecular level.

**Fig 3.**
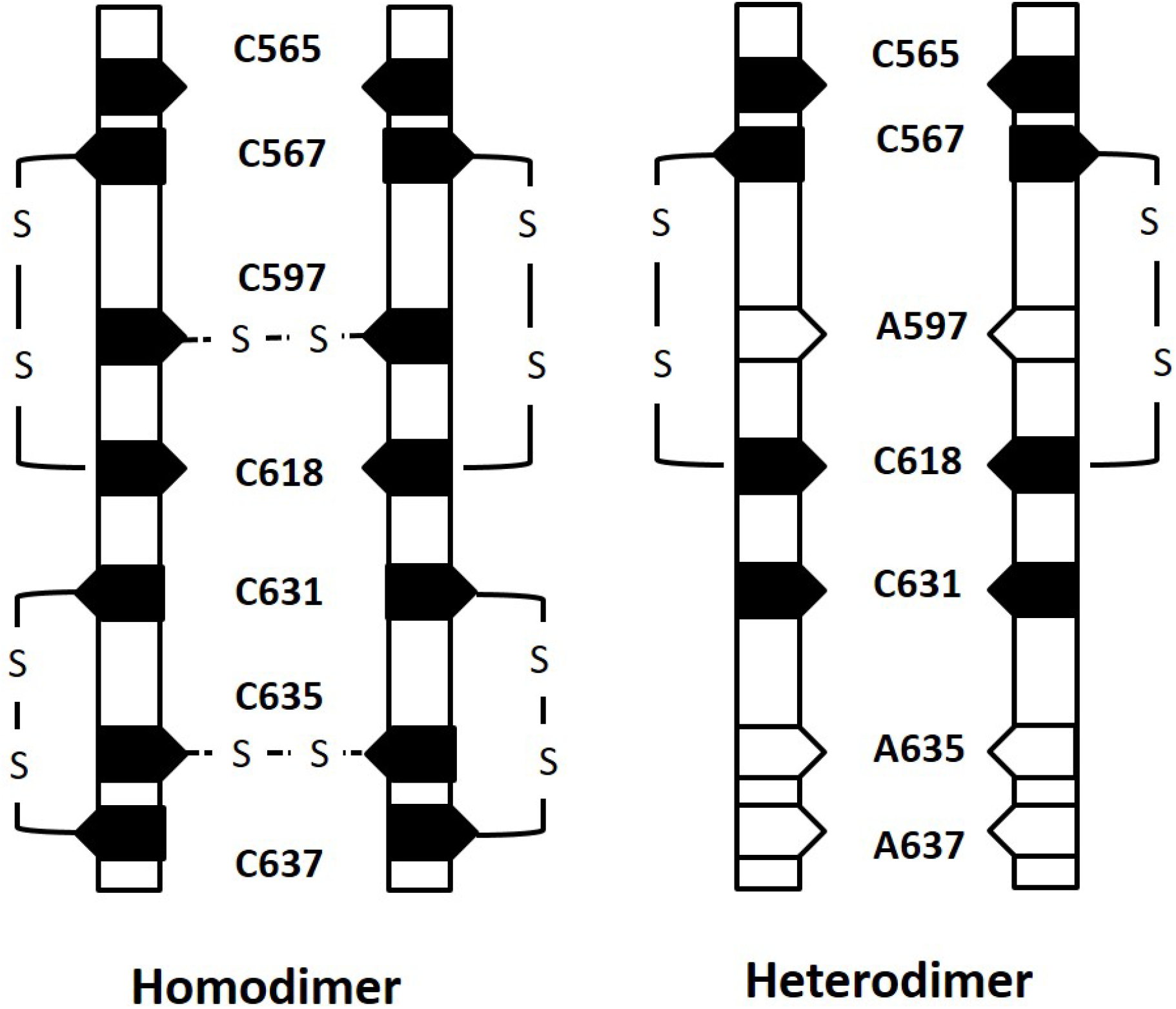
Model explaining role of different cysteines in dimerization. Based on our genetic and mutation data (14, 21) we propose that Cys at 597 and 635 are involved in intermolecular disulphide bond formation which stabilizes the homodimer. Hence, mutations at these two positions disrupt the intermolecular disulfide bonds resulting into loss of Toll-1 homodimer. Thus, Toll-1 monomer resulting from C597 and C635 double mutation will be more favoured to form heterodimer with Toll-9. Further, Cys 565 and 567 favour while Cys at 597, 635 and 637 disfavour heterodimer formation. Since, Cys 565 favours heterodimer formation hence we believe that it might not be involved in intermolecular disulfide bond formation. This indicates critical role played by CRD in homodimer formation.

### 3. Loss of CRD resulted in ability of Toll-9 to recognize dsRNA as ligand

We have established that heterodimerization of toll proteins has evolved due to loss of CRD during the course of evolution. Figure 1 makes it explicit that CRD loss took place at the evolutionary step of non-chordates to chordates transition. This is also the inflection point with respect to functional diversity as organismal complexity increased hereafter tremendously. While insect Toll-proteins are known to bind host proteins as ligands, mammalian Toll proteins form extensive heterodimers and can bind ligands from infecting microbes. Naturally we asked, can Drosophila Toll-9 which lacks CRD and thus resembles mammalian Toll can bind non-protein ligands? We tested LPS, DNA and dsRNA as nonprotein ligands to bind Toll-9. We found positive interaction with dsRNA as seen in the RNA-EMSA (Fig. 3). We confirmed dsRNA binding with recombinant Toll-9 also UV-Crosslinking, mutant analysis and transgenic study (To be published elsewhere).

### 4. Toll-9 monomer has higher affinity for dsRNA than the Toll-9 dimer

One interesting finding about Toll9-dsRNA interaction is that Toll9 can exist in both monomer and dimer forms and also bind the ligand in both forms. This is significant in many ways because Toll-proteins are known to bind ligands as dimers and not as monomers. Our data further makes it clear that dsRNA has higher affinity for the monomer than the dimer. ImageJ analysis confirms that affinity for monomer (66%) is twice to that of the dimer (33%).

There are two possible explanations for this observation; 1) Toll9 has lower propensity to form dimer as it lacks CRD unlike other Toll proteins of Drosophila. Since there are less dimers in the pool hence, less binding with dsRNA. 2) Other possibility is that with dimer formation the surface area available for binding dsRNA is less compared to the monomer. But if this is true then one can safely conclude that the RNA binding surface is towards the interior side which will be less exposed for binding in the dimer form. If RNA binding surface is on the outer side then monomer or dimer will not make a difference with respect to binding affinity. At this moment we believe that explanation 1, the reduced propensity to form dimer due to absence of CRD may be more correct. Other drosophila Toll proteins which have CRD and form dimmers are not known to bind other than proteins as ligands. Thus, we conclude that recognition of dsRNA by Toll9 is a gain of function phenotype resulting from loss of CRD.

### 5. dsRNA recognition by Toll-9, in vivo

dsRNA binding ability of Toll-9 was also confirmed*. in vivo*. as shown in fig.5. here GFP-tagged Toll-9 was expressed in Human HeLa cells and dsRNA was added to the medium later. Since DAPI is an intercalating agent hence can bind dsRNA as well as the DNA (nucleus). One can clearly see DAPI on the cell surface where there is expression of Toll9-GFP (Fig. 5). The cells which are GFP negative also do not show DAPI stained dsRNA on the cell surface. This proves that dsRNA binding to Toll-9 happens in vivo on the cell surface and thus confirms our in vitro data. One can observe increased concentration of Toll-9 and dsRNA complex in dividing cells at the site of cell separation. This may indicate a role for Toll-9 in dividing cells. This is an interesting observation and needs further study (Fig. 5).

**Fig 4.**
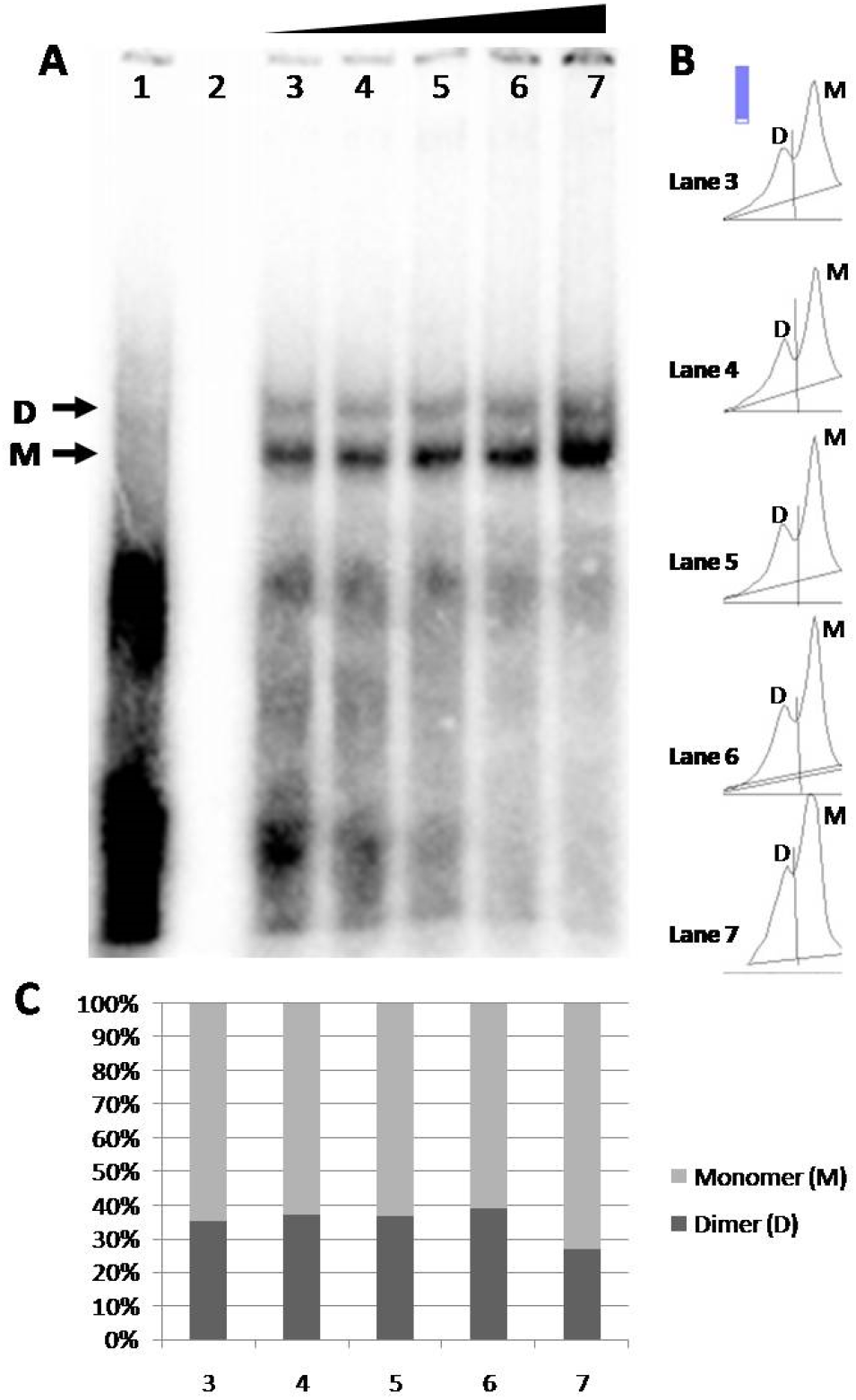
Confirmation of dsRNA binding to Toll9 by RNA-EMSA. (A) A 300 bp long dsRNA was used in EMSA experiments with Toll9 and it formed two complexes probably as monomer (M) and dimer (D). Stronger intensity of dsRNA-monomer complex compared to dsRNA-Monomer complex indicates higher affinity for monomer than dimer. Increasing concentration of dsRNA was used in lanes 3-7. (B&C) ImageJ analysis was performed to quantify monomer and dimer bands (B) which shows that monomer-dsRNA complex was ~67% compared to dimer-dsRNA complex which was around 33%. When dsRNA was increased to 2microgram (Lane7), the monomer complex further increased to 76%. This clearly shows addition of dsRNA favours monomer formation.

**Fig.5.**
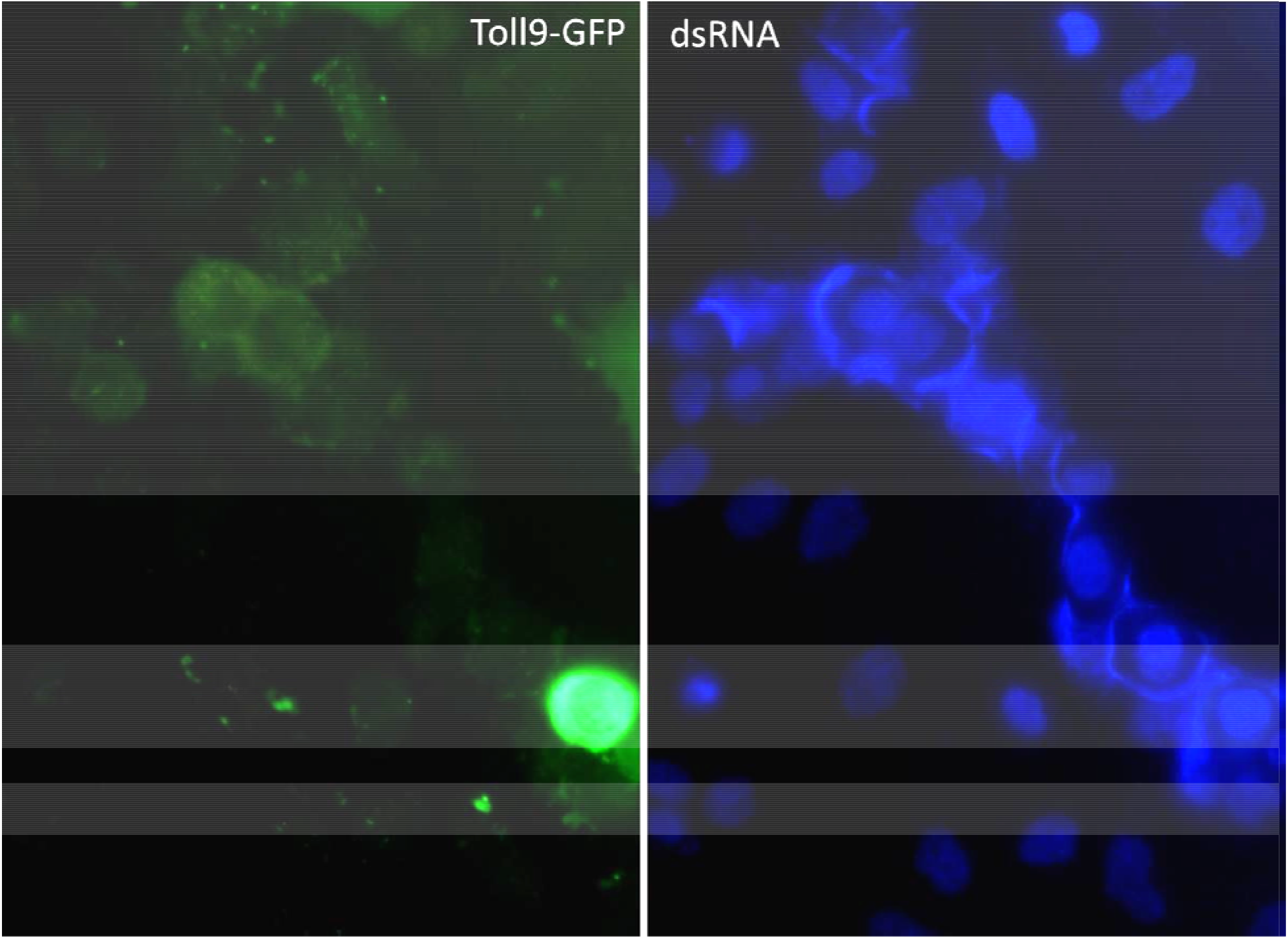
Toll9 binding to dsRNA *in vivo*. EGFP tagged Toll9 was transfected into human HeLa cells and observed under a fluorescent microscope. All GFP positive cells show DAPI staining at the cell surface. Toll9 is a membrane protein and expresses at the cell surface. Binding of dsRNA to Toll9-GFP is confirmed be DAPI staining of the cell surface in GFP positive cells. Cells where GFP is not seen do not show DAPI staining at the cell boundary. This confirms that Toll9 specifically recognizes dsRNA at the cell surface. In other words, dsRNA is bonafide ligand for Drosophila Toll-9.

## Discussion

Toll proteins are involved in plethora of functions and this functional diversity evolved along with organismal complexity. However, the molecular mechanism behind this functional diversity was not known. Here, we have provided an explanation that evolution of heterodimerization in Toll proteins was triggered consequent to the loss of CRD. Thus while other domains of Toll proteins experienced positive selection (LRR, TIR, CRD-TIR) and hence were retained except the LRR-CRD which experienced negative selection and hence was lost during the course of evolution. Loss of LRR-CRD resulted in two gain of function phenotypes (i) evolution of heterodimerization, and (ii) recognition of PAMPs like dsRNA.

Toll proteins with CRD recognize host proteins as ligands while Toll proteins without CRD, as in vertebrates, recognize different types of molecules as ligands including those from pathogens. Hence, we propose that CRD in toll proteins experienced negative selection during evolution, and hence was lost during non-chordates to chordates transition, consequently leading to heterodimerization, recognition of foreign molecules and PAMPs which ultimately resulted in functional diversity.

It is evident from this study that dimer formation in Toll proteins is regulated primarily by LRR. However, CRD decides formation of homo/hetero-dimers. CRD may impart structural rigidity in insect Tolls and hence the flexibility to form heterodimers with other Toll paralogs would be less (Fig. 3). This is expected as CRDs of different Toll paralogs are not present at the same position and hence can’t align symmetrically in order to form disulfide bridges. This would limit their ability to form heterodimers. On the other hand, mammalian TLRs lack CRD and hence dimer formation is regulated only by LRRs which makes them more flexible to interact with LRRs of other Toll proteins leading to formation of different heterodimers. Thus, we propose that heterodimerization of toll proteins is an evolved property.

One can also not ignore the fact that mammalian Toll proteins directly recognize the PAMPs while insect Toll proteins don’t interact with PAMPs. This is the first report where a PAMP, dsRNA, has been shown to bind an insect Toll-protein. Till date Drosophila Toll proteins have only been shown to bind host proteins as ligands not even foreign protein. In case of infection molecular pathway is triggered such that it activates appropriate host protein to bind as ligand to the cognate Toll protein to initiate downstream signaling. We propose that this limitation in insect toll proteins is imparted by the presence of CRD. It is also tempting to propose that Toll proteins originated as regulators of developmental processes and later on evolved pathogen recognition. We believe that loss of CRD played an important role in this transition.

## Acknowledgements

Authors declare no conflict of interest. NM is recipient of IYBA grant from Department of Biotechnology. PR is a recipient of Post-Doctoral Fellowship from UGC, Govt. of India. NM conceived the project and designed the experiments. PR, SKS and NM performed the experiments. NM wrote the manuscript.

## Materials and Methods

### PCR amplification of toll-1 and toll-9 ectodomains

The 2.4 kb etoll and 1.5 kb etoll-9 were PCR amplified from cDNA pool of wild D. melanogaster with Nde1 and BamH1 restriction sites at their 5’ and 3’ ends respectively. Primers used were 5’-AGTCGACTAAGGCCGCTTC (forward primer, FP) and 5’-CACAGCAAGGGCTATGGAACA (reverse primer, RP) for etoll; 5’-GCCTGTTCCTTGGAAACGTA (forward primer, FP) and 5’-GTTGGACAGCATGTTGATGG (reverse primer, RP) for etoll-9. These primers were allowed to anneal to the template at 58° C for 1 minute. Further, amplified constructs of etoll and etoll-9 were cloned in the MCS of pGADT7-AD and pGBKT7-BD vector respectively using the same restriction enzymes. Restriction digestion and nucleotide sequencing later confirmed positive clones.

### Generation of CRD-deleted constructs of toll ectodomain

Overlap extension PCR was used for deletion of 200 bp (approx.) N-terminal CRD from etoll (supplementary figure). For amplification of distal fragment (1.7 kb approx.), primers used were 5’-TCCCATATGATGAGTCGACTAAAGGCCGC (forward primer, FP) and 5’-TAGGTTCGCACATGGACCAGGGGATTATCGTTGAG (reverse primer, R1). Primers used for amplification of proximal fragment (0.5 kb approx.) were 5’-ATAATCCCCTGGTCCATGTGCGAACCTACGACAAAGC (forward primer, F1) and 5’-TTAGGATCCCACAGCAAGGGCTATGAACA (reverse primer, RP). Standard PCR was used and annealing of primers in both the cases were done at 58 °C for 1 minute.

Full length ΔCRD etoll was generated by integrating the distal fragment with the proximal fragment. This OE PCR was carried out in two steps: in the first step, distal and proximal fragments were allowed to overlap their common sequences followed by extension. Reactions of both fragments were pooled and was run as follows: initial 3 min. at 95°C, 15 cycles of amplification (30 sec at 95°C, 1 min at 58°C and 2 min at 72°C) and a final extension for 10 min at 72°C. In the second step, full length ΔCRD etoll was subject to amplification using primers FP and RP with annealing step at 58°C for 1 min 30 sec. and further ligated into pcDNA 3.1 V5/His TOPO vector using pcDNA 3.1 V5/His TOPO TA cloning kit (Invitrogen). The ΔCRD etoll was then transferred to pGADT7-AD vector using Nde1 and BamH1 sites. It was then confirmed by restriction digestion and sequencing.

### Site directed mutagenesis

Cys to Ala substitution mutations were introduced at specific sites in the ectodomain of wild type toll by using QuickChange II XL site-directed mutagenesis kit (Stratagene) as per the manufacturer’s protocol. Primers mentioned in the supplementary table were used for incorporating single or double mutations (supplementary table). Appropriate mutants were confirmed by sequencing (supplementary data).

### Yeast-two hybrid assay

For verification of protein-protein interactions Gal4 Two-Hybrid system was used. All the mutants of etoll were cloned separately in pGADT7-AD vector (Clontech) and wild type etoll-9 was cloned in pGBKT7-BD vector (Clontech). Each mutant etoll-AD was transformed in Y187 strain followed by selection on synthetic defined (SD) −Leu dropout plates whereas etoll-9-BD was transformed in AH109 strain and plated on SD −Trp dropout plate. For mating of Y187 (pAD-mutant etoll) with AH109 (pBD-etoll-9), a colony of each were incubated at 30°C for 1-2 days. This step was performed separately for each mutant of etoll with wt etoll-9. For positive control, Y187 (pAD-T-ag) and AH109 (pBD-p53) were used and Y187 (pAD-empty vector) and AH109 (pBD-empty vector) as negative control. Further, diploids were selected on SD −Leu—Trp dropout plates. These double dropout plates allowed growth of yeast cells with both pAD and pBD. Transformants were further streaked onto SD −Leu −Trp −His −Ade plates, which were incubated at 30°C for 2-4 days to examine the growth. Growth of yeast cells on the quadruple dropout plates were a result of activation of reporter gene due to interaction of the two fusion proteins.

### Cell transfection and Imaging

Full length Toll-9 was cloned into pEGFP-N1 vector where GFP is downstream to Toll9 cDNA. 5 microgram of This plasmid was transfected into HeLa cells using cellfectin (Invitrogen) reagent. 24 hours post transfections the cells were washed and fresh media added. 6 hours later to this dsRNA was added into the media. 6 hours post dsRNA addition the cells were fixed and observed under a fluorescent microscope for imaging.

### Expression and purification of Toll-9

pET-Duet-Toll-9 full length clone was transformed into BL21 (DE3) (Novagen) for the expression. Protein was induced using 0.1mM IPTG for a period of 3hrs at 37 °C. Total protein extracts were prepared by using B-PER^®^ Bacterial Protein Extraction Reagent (Thermo Scientific) as per manual. The protein was resolubilized from inclusion bodies by using Inclusion Body Solubilization Reagent (Thermo Scientific) following manufacturer’s protocol. The protein mixture was dialyzed against one liter of 6M urea overnight using SnakeSkin Dialysis Tubing, 3.5K MWCO (Thermo Scientific) at 4 °C. 250mL of 25mM Tris-HCl (pH 7.5) was added to the beaker every 6 hours until the final volume reached to 3 liters, later replaced with 2L of 25mM Tris-HCl (pH 7.5) and 150mM NaCl overnight at 4 °C. For purification, refolded protein mixture was loaded on Ni-NTA beads (Qiagen) as 1ml per 2 liter culture, pre-washed with 25 mM Tris-HCl pH 8.0. To reduce noise, beads were washed with washing buffer (25 mM Tris-HCl, pH 8.0) with a gradient concentration of imidazole (10-50 mM). Finally, beads were treated with elution buffer (25 mM Tris-HCl, pH 8.0, imidazole 250 mM). The eluted fraction was dialyzed against the buffer 25 mM Tris-HCl, pH 8.0, 5% glycerol, 1mM DTT flash frozen and stored at −80□°C.

### dsRNA preparation

A 150bp of dsRNA was synthesized by using empty pCR^®^2.1-TOPO vector as a template which has both SP6 and T7 promoters. M13 primers were used for the amplification by PCR. For the synthesis of 300bp dsRNA, a region of *Bombyx mori intersex* gene was amplified by specific primers flanked by SP6 and T7 promoters in opposite directions. Sense and antisense RNA were transcribed by using Megascript T7 and SP6 *in vitro* transcription kit as per owner’s instructions (Thermo Fischer Scientific). To remove contaminating PCR products after transcription, DNase I digestion was given. The sense and antisense RNA were precipitated by LiCl at −70 °C for 1hr. After centrifugation, the RNA Pellet was reconstituted in water and concentration was estimated by NanoDrop 2000 UV-Vis Spectrophotometer (Thermo Fisher Scientific).

### dsRNA labeling

Various dsRNA probes were generated by end labeling reaction. Briefly, 12.5 pmol of dsRNA, 1μl of (12 U/μl) T4-polynucleotide kinase (PNK), 2.5μl of 10x T4 PNK buffer, 4μl of [γ-^32^P] ATP (sp. Activity 3200 Ci/mmol, 10mCi/mL) were mixed and the volume was made to 25μl with DEPC-treated water. The reaction mixture was incubated at 37°C for 30 min. and terminated at 70°C for 10 min. The dsRNA probe was purified by using G-25 MicroSpin column (GE Healthcare). Geiger–Muller counter was used to detect the transfer of radio-label.

### RNA Electrophoretic Mobility Shift Assay

For RNA-EMSA studies, radiolabeled probe (30,000 cpm) was incubated with varying concentrations of recombinant proteins in 1X binding buffer (10 mM HEPES (pH 7), 50 mM KCl, 1 mM EDTA, 1 mM DTT, 10% glycerol and 0.5% Triton X-100) along with 1μg of poly-dI/dC and water to make the final reaction volume to 30μl. The reaction mixture was incubated at room temperature for 30 min. For the competitive assay, the various amounts of cold competitor were incubated in the binding reaction mixture, 30 min before the addition of the probe (labeled oligo). The samples were resolved on to a 6% native polyacrylamide gel (29:1) at 200V in cold room for 2-3 hrs with 1X TBE buffer. The gel was pre-ran at 100V for 30 min at 4 °C. Post run, gels were transferred to Whatman paper 3, dried and visualized using a PhosphorImager (Typhoon FLA 9500, GE).

---

**Fig S1.**
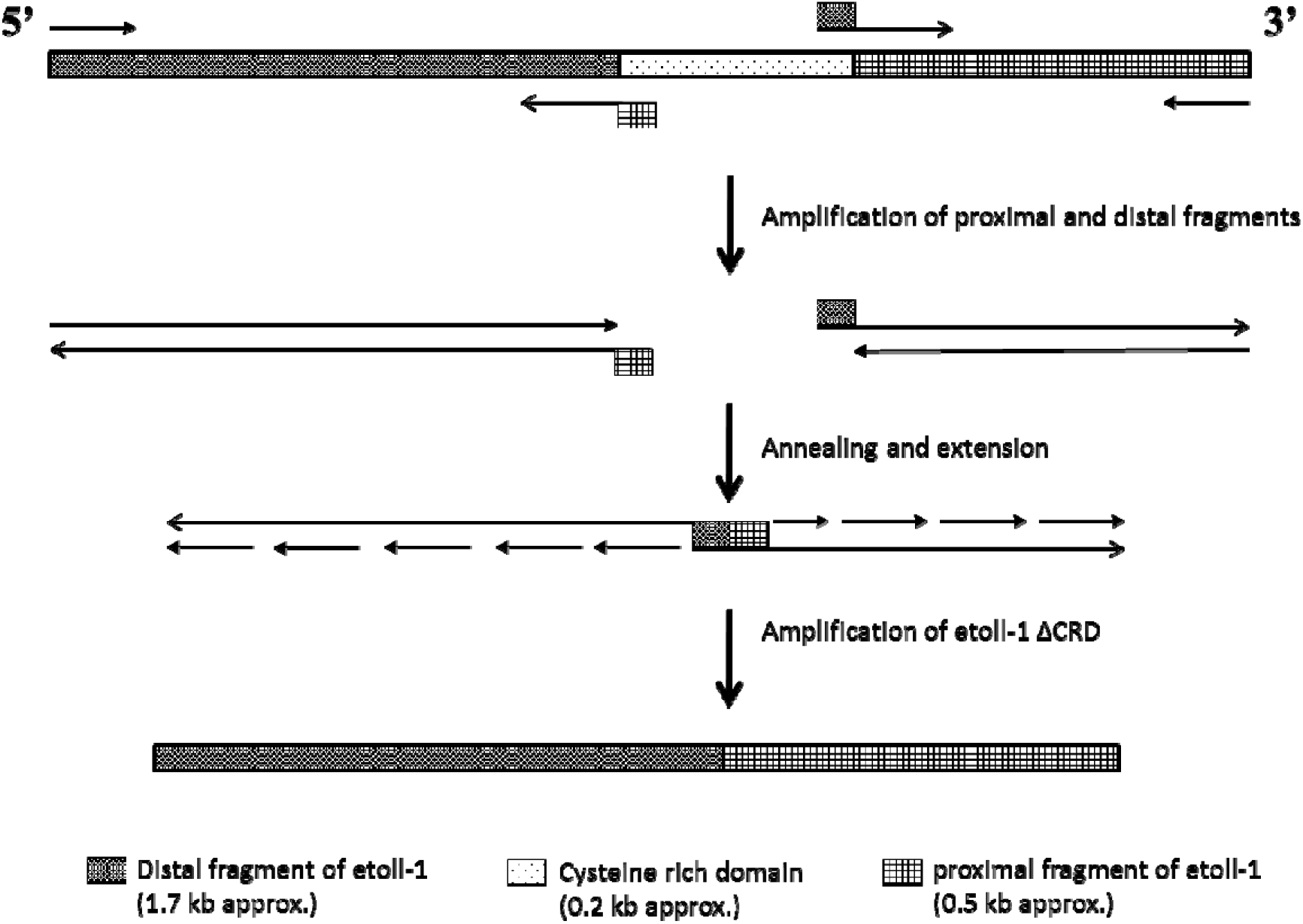
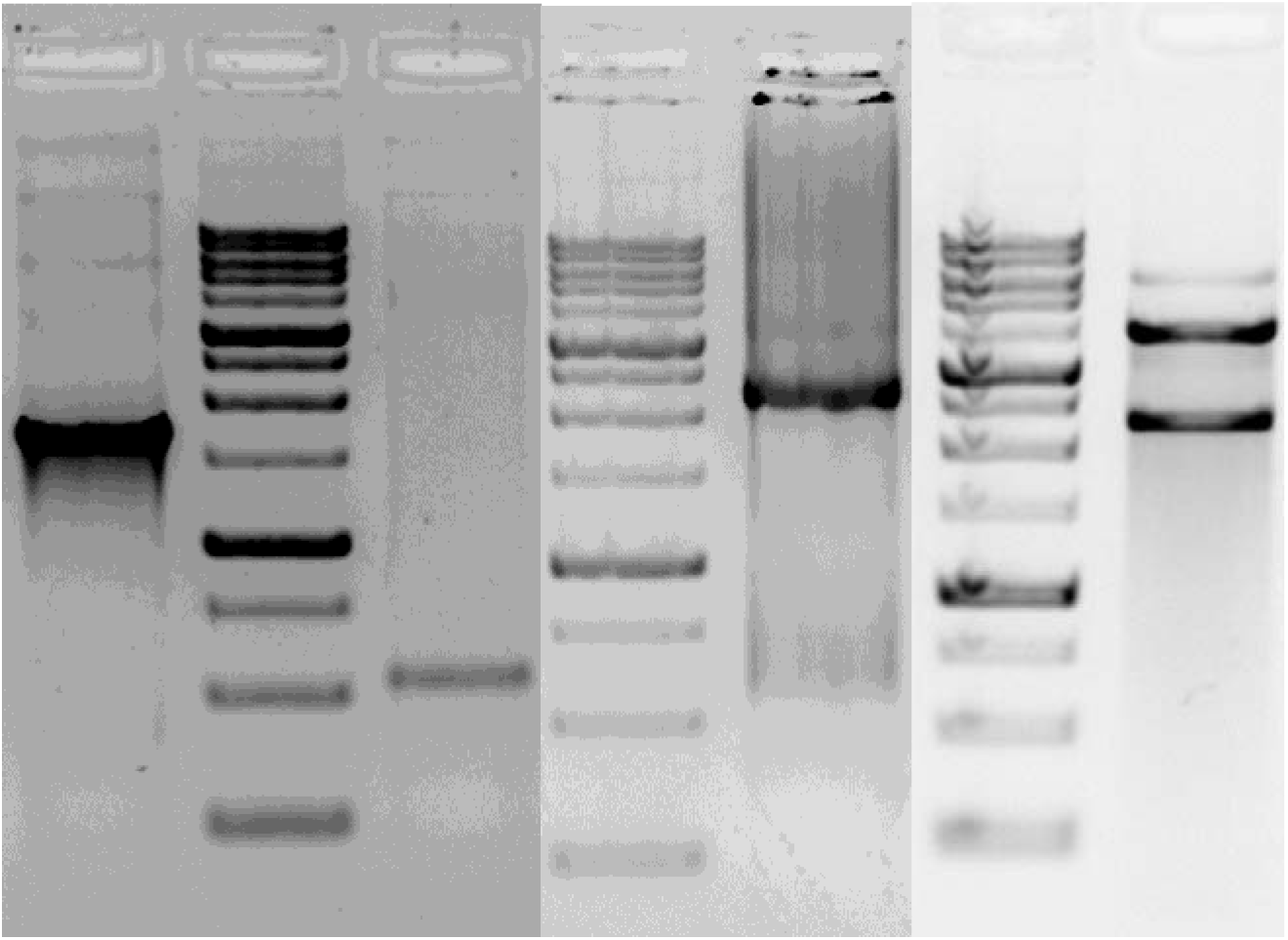
Generation of CRD deletion construct of Toll-1 ectodomain. Agarose gel electrophoresis showing amplification of distal fragment of toll-1 ectodomain (etoll-1) (lane 1) and proximal fragment of toll ectodomain (lane 2). Both fragments were overlapped, extended, and amplified to generate etoll-1 ΔCRD (lane 3) which was further check through digestion with NdeI and BamHI (lane 4).

**Fig S2.**
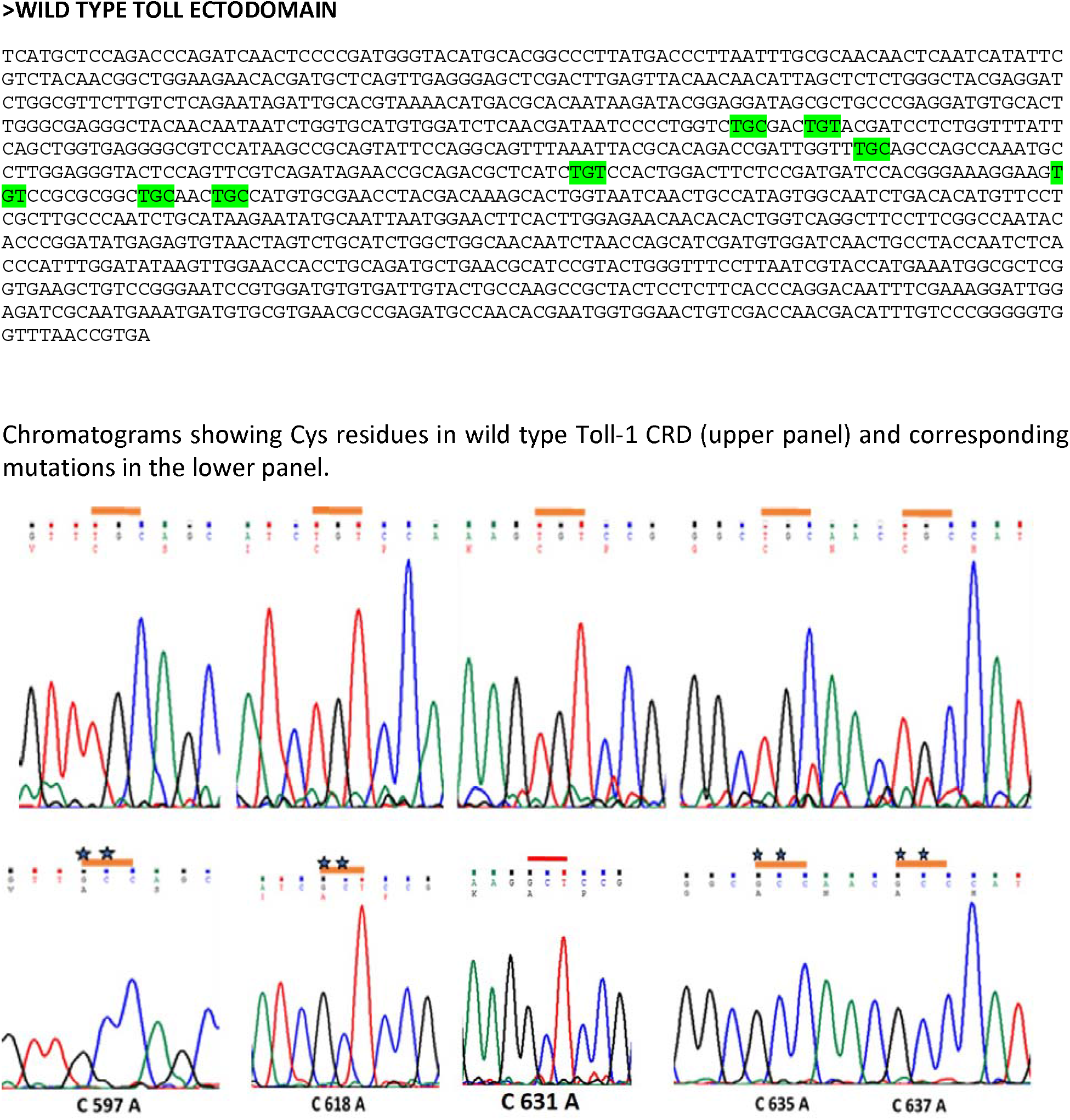
Sequence of wild-type and different site-directed mutants (SDM) along with their chromatograms.

**Fig S3.**
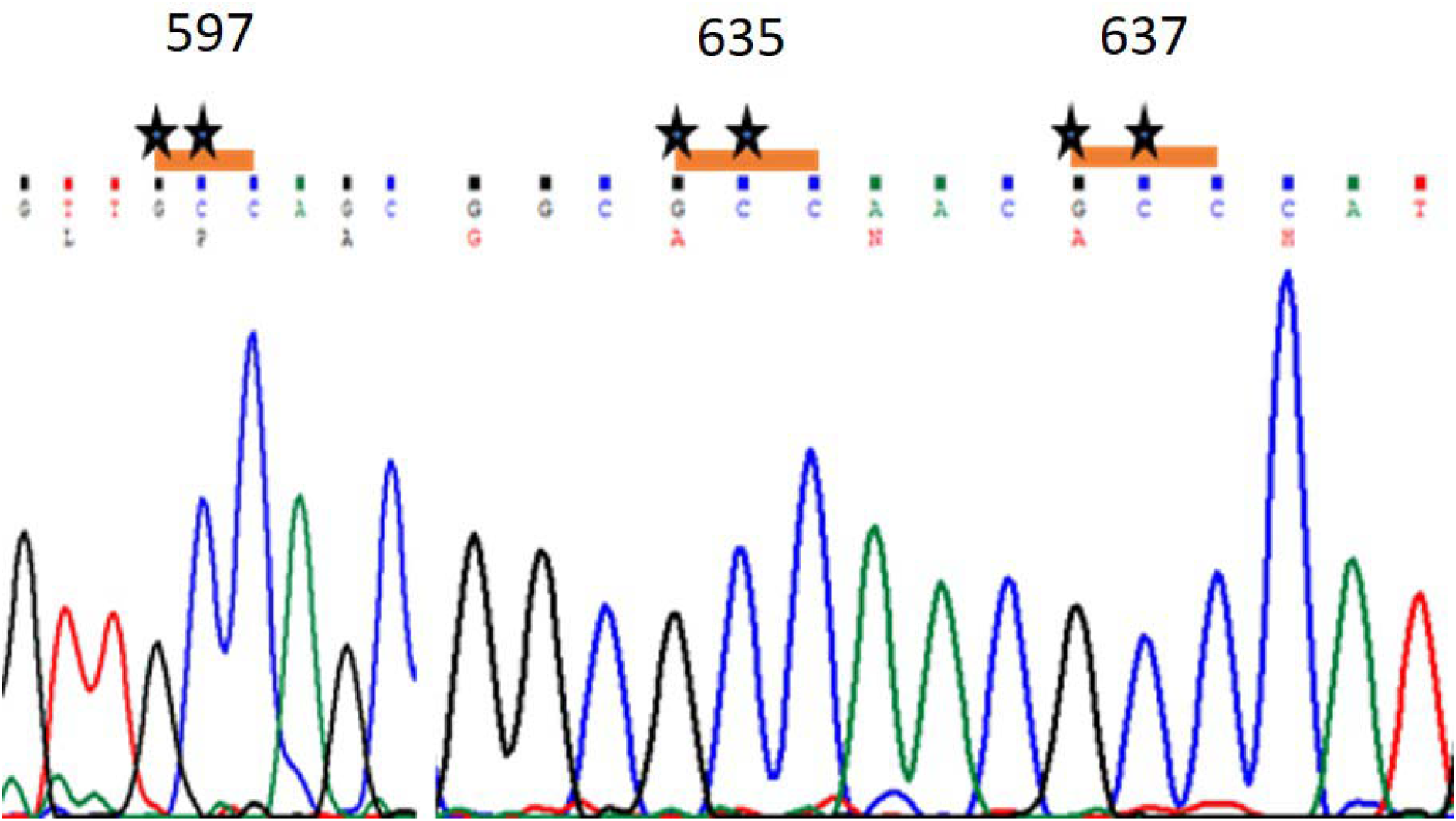
Chromatogram of SDM25 mutant. Chromatogram shows Cys mutations at positions 597, 635 and 637 in a single construct named as SDM 25. This construct shows interaction with toll9 indicating that Cys at these positions inhibit heterodimer formation while a mutation here favours the same.

**Fig S4.**
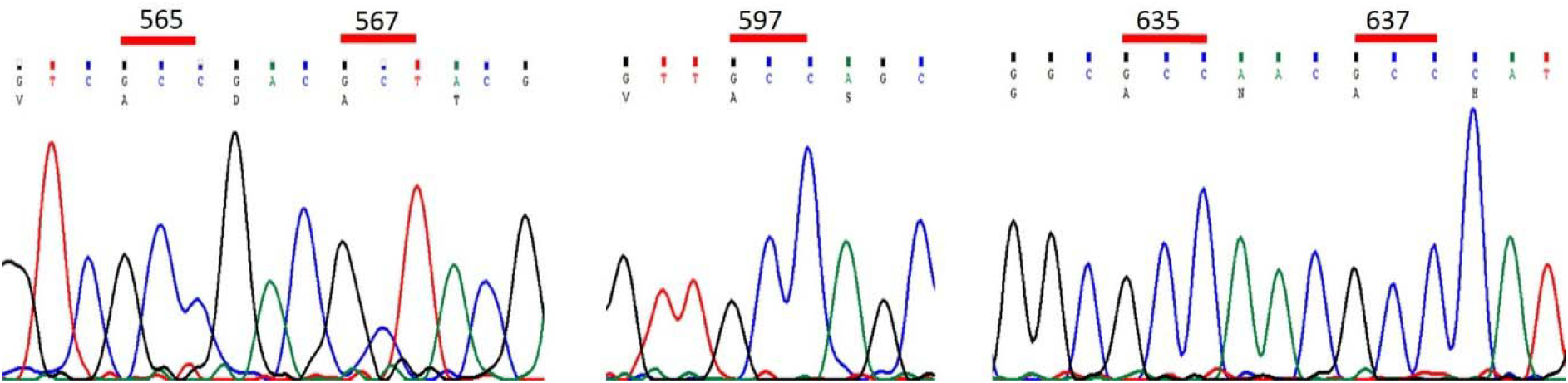
Chromatogram of SDM 125. Chromatogram shows Cys mutations at positions 565, 567, 597, 635 and 637 in a single construct named as SDM125.

## References

1. K. V. Anderson, G. Jurgens, C. Nusslein-Volhard, Establishment of dorsal-ventral polarity in the *Drosophila* embryo: genetic studies on the role of the Toll gene product. Cell. 42, 779–789 (1985).

2. S. Akira, K. Takeda, Toll-like receptor signalling. Nat. Rev. Immunol. 4, 499–511 (2004).

3. R. Medzhitov, P. Preston-Hurlburt, C. A. Janeway Jr, A human homologue of the *Drosophila* Toll protein signals activation of adaptive immunity. Nature. 388, 394–397 (1997).

4. N. Silverman, T. Maniatis, NF-kappaB signaling pathways in mammalian and insect innate immunity. Genes Dev. 15, 2321–2342 (2001).

5. J. C. Sullivan, D. Kalaitzidis, T. D. Gilmore, J. R. Finnerty, Rel homology domain-containing transcription factors in the cnidarian Nematostella vectensis. Dev. Genes Evol. 217, 63–72 (2007).

6. N. H. Putnam, M. Srivastava, U. Hellsten, B. Dirks, J. Chapman, et al., Sea anemone genome reveals ancestral eumetazoan gene repertoire and genomic organization. Science. 317, 86–94 (2007).

7. M. Wiens, M. Korzhev, S. Perovic-Ottstadt, B. Luthringer, D. Brandt, et al., Toll-like receptors are part of the innate immune defense system of sponges (demospongiae: Porifera). Mol. Biol. Evol. 24, 792–804 (2007).

8. B. Kobe, J. Deisenhofer, Proteins with leucine-rich repeats. Curr. Opin. Struct. Biol. 5, 409–416 (1995).

9. P. Enkhbayar, M. Kamiya, M. Osaki, T. Matsumoto, N. Matsushima, Structural principles of leucine-rich repeat (LRR) proteins. Proteins. 54, 394–403 (2004).

10. B. Kobe, A. V. Kajava, The leucine-rich repeat as a protein recognition motif. Curr. Opin. Struct. Biol. 11, 725–732 (2001).

11. E. Latz, A. Verma, A. Visintin, M. Gong, C. M. Sirois, Ligand-induced conformational changes allosterically activate Toll-like receptor 9. Nat. Immunol. 8, 772–779 (2007).

12. A. Ozinsky, D. M. Underhill, J. D. Fontenot, A. M. Hajjar, The repertoire for pattern recognition of pathogens by the innate immune system is defined by cooperation between toll-like receptors. Proc. Natl. Acad. Sci. U. S. A., 97, 13766–13771 (2000).

13. M. Triantafilou, F. G. Gamper, R. M. Haston, M. A. Mouratis, S. Morath, Membrane sorting of toll-like receptor (TLR)-2/6 and TLR2/1 heterodimers at the cell surface determines heterotypic associations with CD36 and intracellular targeting. J. Biol. Chem. 281, 31002–31011 (2006).

14. C. Parthier, M. Stelter, C. Ursel, U. Fandrich, H. Lilie, Structure of the Toll-Spatzle complex, a molecular hub in *Drosophila* development and innate immunity. Proc. Natl. Acad, Sci. U. S. A. 111, 6281–6286 (2014).

15. J. L. Imler, L. Zheng, Biology of Toll receptors: lessons from insects and mammals. J Leukoc Biol, 75, 18–26 (2004).

16. F. Leulier, B. Lemaitre, Toll-like receptors--taking an evolutionary approach. Nat Rev Genet, 9, 165–178 (2008).

17. Z. Kambris, J. A. Hoffmann, J. L. Imler, M. Capovilla, Tissue and stage-specific expression of the Tolls in *Drosophila* embryos. Gene Expr Patterns, 2, 311–317 (2002).

18. M. S. Jin, J. O. Lee, Structures of the toll-like receptor family and its ligand complexes. Immunity, 29, 182–191 (2008).

19. A. N. Weber, S. Tauszig-Delamasure, J. A. Hoffmann, E. Lelievre, H. Gascan et al., Binding of the *Drosophila* cytokine Spatzle to Toll is direct and establishes signaling. Nat. Immunol, 4, 794–800 (2003).

20. A. N. Weber, M. C. Moncrieffe, M. Gangloff, J. L. Imler, N. J. Gay, Ligand-receptor and receptor-receptor interactions act in concert to activate signaling in the *Drosophila* toll pathway. J Biol Chem. 280, 22793–22799 (2005).

21. I. Botos, D. M. Segal, D. R. Davies, The structural biology of Toll-like receptors. Structure. 19, 447–459 (2011).

22. R. D. Bies, C. T. Caskey, R. Fenwick, An intact cysteine-rich domain is required for dystrophin function. Journal of Clinical Investigation. 90, 666–672 (1992).

23. M. Branschadel, A. Aird, A. Zappe, C. Tietz, A. Krippner-Heidenreich, et al., Dual function of cysteine rich domain (CRD) 1 of TNF receptor type 1: conformational stabilization of CRD2 and control of receptor responsiveness. Cell Signal. 22, 404–414 (2010).

24. K. Yu, K. H. Kang, P. Heine, U. Pyati, S. Srinivasan, et al., Cysteine repeat domains and adjacent sequences determine distinct bone morphogenetic protein modulatory activities of the *Drosophila* Sog protein. Genetics. 166, 1323–1336 (2004).

25. M. Gangloff, A. Murali, J. Xiong, C. J. Arnot, A.N. Weber, et al., Structural insight into the mechanism of activation of the Toll receptor by the dimeric ligand Spatzle. J Biol Chem. 283, 14629–14635 (2008).

