## Supplemental figures for "Loss of Cysteine Rich domain was critical for evolution of heterodimerization in Toll proteins"

**Supplemental material**

**Materials and Methods**

*PCR amplification of toll and toll-9 ectodomain*

The 2.4 kb e-toll (extracellular domain of Toll1) and 1.5 kb e-toll-9 (extracellular domain of Toll-9) were PCR amplified from cDNA pool of CantonS D. melanogaster with Nde1 and BamH1 restriction sites at their 5’ and 3’ ends respectively. Primers used were 5’-AGTCGACTAAGGCCGCTTC (forward primer, FP) and 5’- CACAGCAAGGGCTATGGAACA (reverse primer, RP) for etoll; 5’- GCCTGTTCCTTGGAAACGTA (forward primer, FP) and 5’- GTTGGACAGCATGTTGATGG (reverse primer, RP) for etoll-9. These primers were allowed to anneal to the template at 58° C for 1 minute. Further, amplified constructs of etoll and etoll-9 were cloned in the MCS of pGADT7-AD and pGBKT7-BD vector respectively using the same restriction enzymes. Restriction digestion and nucleotide sequencing later confirmed positive clones.

*UV Cross-linking assay*

For cross-linking assays, RNA-Protein Complexes were prepared as described in EMSA. Samples were exposed to UV rays in Stratalinker ((Stratagene 2400), with 999.9 mJ/cm2 energy on ice for 1 min, three times intermittently. Samples were resolved on 10% SDS-PAGE under reducing conditions. Drying and visualization of the gel were done using a PhosphorImager (Typhoon FLA 9500, GE).

**Supplementary Figures S1-S4.**

**Fig S1. Generation of CRD deletion construct of Toll-1 ectodomain.**

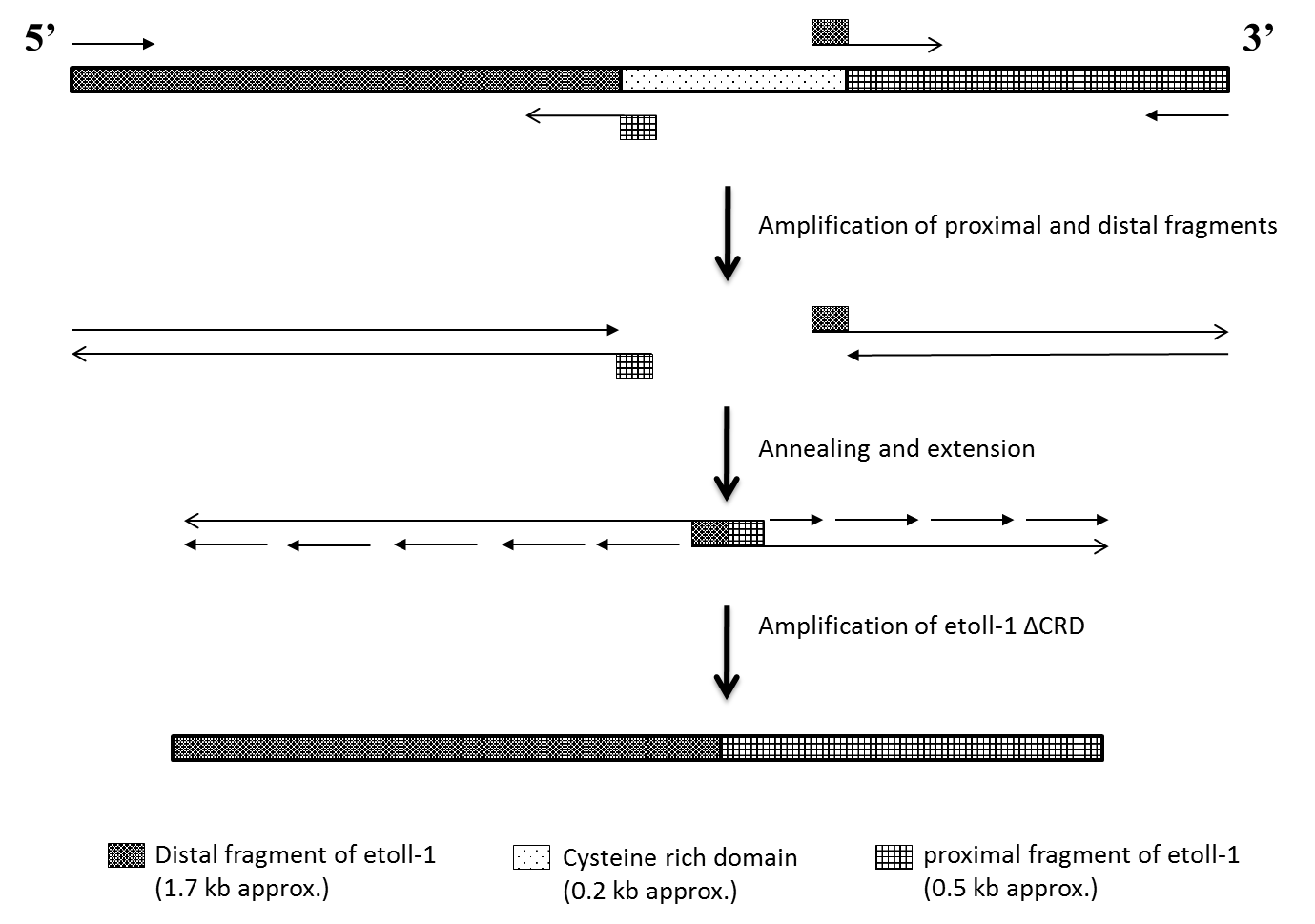

**1	M	2	M	3	M	4**

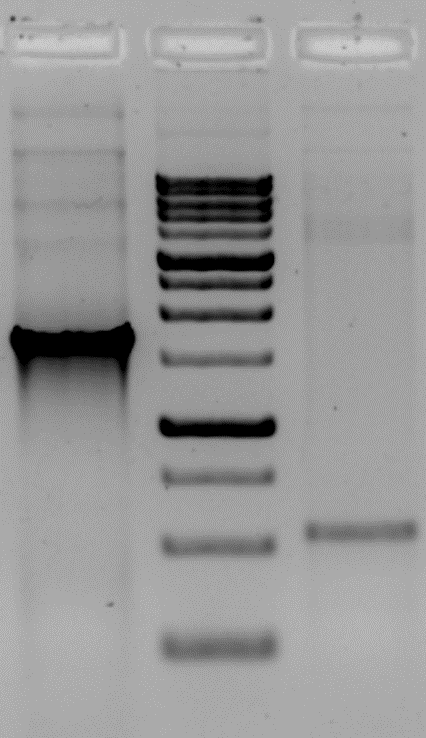

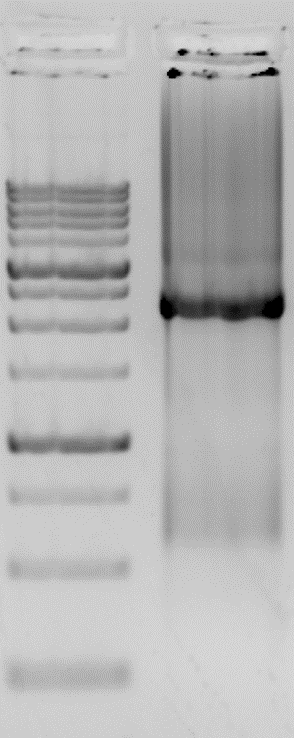

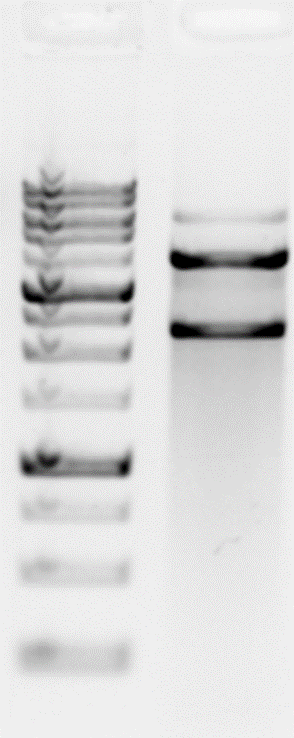

From left to right-

Agarose gel electrophoresis showing amplification of distal fragment of toll-1 ectodomain (etoll-1) (lane 1) and proximal fragment of toll ectodomain (lane 2). Both fragments were overlapped, extended, and amplified to generate etoll-1 ΔCRD (lane 3) which was further check through digestion with NdeI and BamHI (lane 4).

**Fig S2.Sequence of wild-type and different site-directed mutants (SDM) along with their chromatograms.**

**>WILD TYPE TOLL ECTODOMAIN**

TCATGCTCCAGACCCAGATCAACTCCCCGATGGGTACATGCACGGCCCTTATGACCCTTAATTTGCGCAACAACTCAATCATATTCGTCTACAACGGCTGGAAGAACACGATGCTCAGTTGAGGGAGCTCGACTTGAGTTACAACAACATTAGCTCTCTGGGCTACGAGGATCTGGCGTTCTTGTCTCAGAATAGATTGCACGTAAAACATGACGCACAATAAGATACGGAGGATAGCGCTGCCCGAGGATGTGCACTTGGGCGAGGGCTACAACAATAATCTGGTGCATGTGGATCTCAACGATAATCCCCTGGTCTGCGACTGTACGATCCTCTGGTTTATTCAGCTGGTGAGGGGCGTCCATAAGCCGCAGTATTCCAGGCAGTTTAAATTACGCACAGACCGATTGGTTTGCAGCCAGCCAAATGCCTTGGAGGGTACTCCAGTTCGTCAGATAGAACCGCAGACGCTCATCTGTCCACTGGACTTCTCCGATGATCCACGGGAAAGGAAGTGTCCGCGCGGCTGCAACTGCCATGTGCGAACCTACGACAAAGCACTGGTAATCAACTGCCATAGTGGCAATCTGACACATGTTCCTCGCTTGCCCAATCTGCATAAGAATATGCAATTAATGGAACTTCACTTGGAGAACAACACACTGGTCAGGCTTCCTTCGGCCAATACACCCGGATATGAGAGTGTAACTAGTCTGCATCTGGCTGGCAACAATCTAACCAGCATCGATGTGGATCAACTGCCTACCAATCTCACCCATTTGGATATAAGTTGGAACCACCTGCAGATGCTGAACGCATCCGTACTGGGTTTCCTTAATCGTACCATGAAATGGCGCTCGGTGAAGCTGTCCGGGAATCCGTGGATGTGTGATTGTACTGCCAAGCCGCTACTCCTCTTCACCCAGGACAATTTCGAAAGGATTGGAGATCGCAATGAAATGATGTGCGTGAACGCCGAGATGCCAACACGAATGGTGGAACTGTCGACCAACGACATTTGTCCCGGGGGTGGTTTAACCGTGA

Chromatograms showing Cys residues in wild type Toll-1 CRD (upper panel) and corresponding mutations in the lower panel.

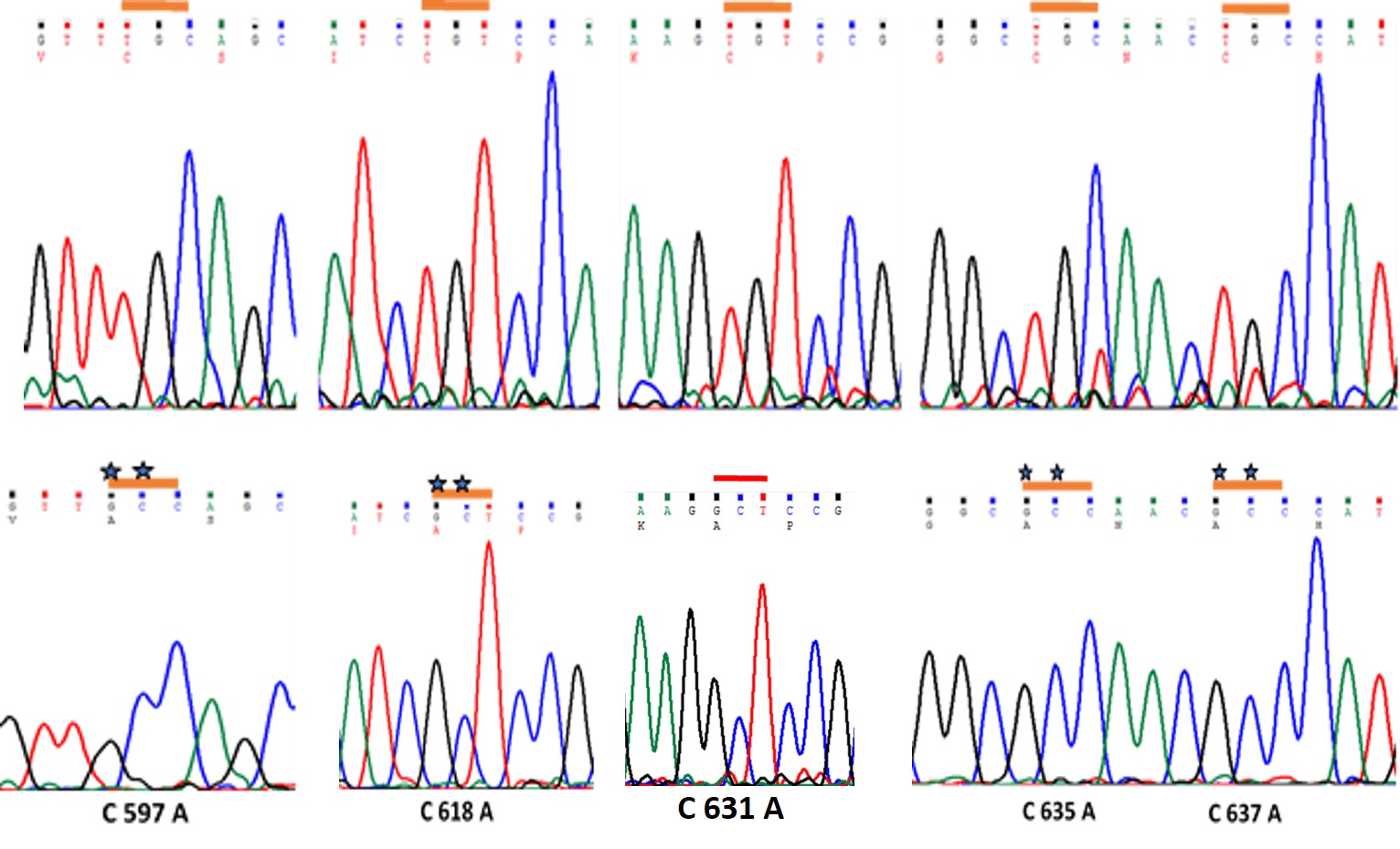

**Fig S3. Chromatogram of SDM25 mutant.**

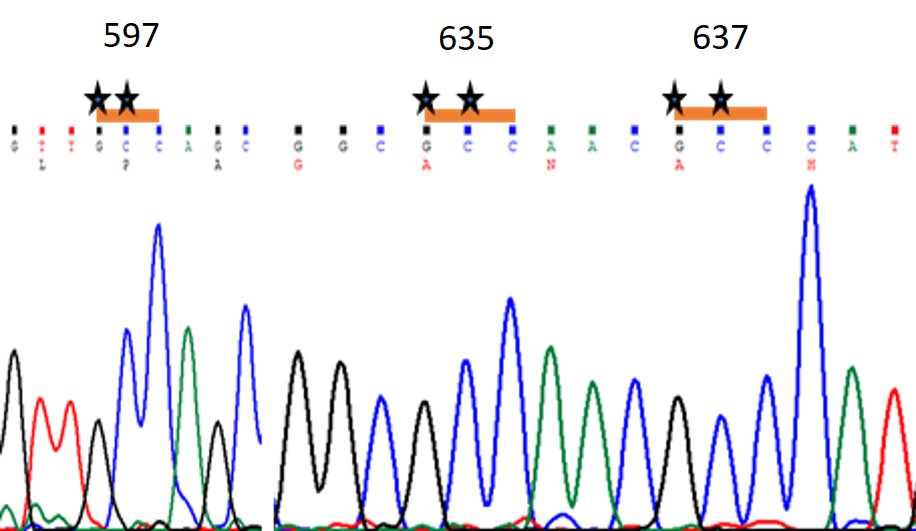

Chromatogram shows Cys mutations at positions 597, 635 and 637 in a single construct named as SDM 25. This construct shows interaction with toll9 indicating that Cys at these positions inhibit heterodimer formation while a mutation here favours the same.

**Fig S4. Chromatogram of SDM 125.**

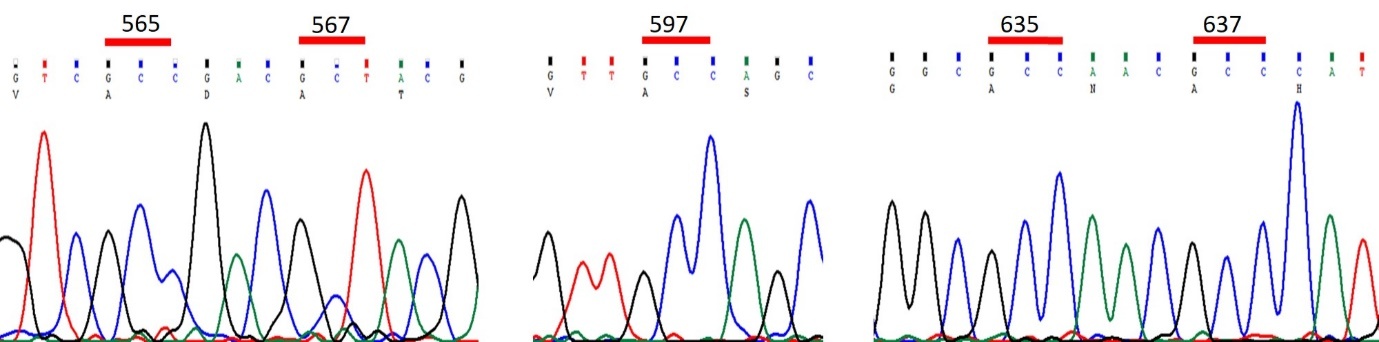

Chromatogram shows Cys mutations at positions 565, 567, 597, 635 and 637 in a single construct named as SDM125.
